# Increased offspring size and reduced gestation length in an ectothermic vertebrate under a worst-case climate change scenario

**DOI:** 10.1101/2023.11.25.568677

**Authors:** David L Hubert, Ehren J Bentz, Robert T Mason

## Abstract

As global temperatures continue to increase, understanding the impacts of warming environments is increasingly relevant. Temperature is especially relevant for ectothermic organisms which depend upon consistent and predictable annual temperature cycles for reproduction and development. However, additional research is required in this area to elucidate the potential impacts of climate change on future generations. To understand how projected increases in environmental temperatures may impact reproductive outcomes within natural populations of ectothermic vertebrates, we manipulated minimum ambient temperatures during gestation in Red-sided garter snakes (*Thamnophis sirtalis parietalis*). Wild snakes were collected in the Interlake region of Manitoba, Canada during their spring mating season and allowed to mate in controlled conditions. For the duration of gestation, mated females were placed into one of two ambient thermal conditions: temperatures emulating those found in the species’ natural habitat or temperatures with a consistent 5 °C increase to match end-of-century climate change projections. We recorded observations for each litter and all neonates resulting from controlled mating trials. We observed no difference in litter sizes or birth rates between thermal conditions. However, we observed a significant reduction in gestation length and significant increase to neonate body mass and body condition associated with increased ambient temperatures. These results suggest that increased minimum temperatures during gestation may confer reproductive benefits for the northern populations of this species even under the most extreme current modeled warming predictions. We discuss the broader implications of this effect, including possible negative ecological outcomes.

## 1. Introduction

In a time of rapidly changing environmental temperatures, it is increasingly important to elucidate how altered thermal conditions impact the coordination of important life history stages and essential events such as mating, gestation, and foraging. Because reproductive success is a defining feature of individual fitness, factors that greatly impact reproduction among individuals within a population have the potential to dramatically affect the long-term success and survival of that population (Brommer et al, 2002; Sherman and Runge, 2002; Rabaiotti et al., 2023). Ectotherms are influenced particularly strongly by environmental conditions, especially temperature, which is frequently an important cue used to coordinate behavior and physiological processes essential to reproductive success including courtship behavior and successful incubation or gestation (Huey and Stevenson, 1979; Huey, 1982; Miles et al., 2000; Lourdais et al., 2013). Ectotherms from terrestrial ecosystems demonstrate existing plastic physiological and behavioral mechanisms to maintain homeostasis during large seasonal and acute temperature fluctuations (Huey et al., 2001; Gunderson et al., 2017; Noer et al., 2022). These extreme temperature fluctuations have been increasingly exacerbated by recent climate change (Masson-Delmotte et al., 2021). Therefore, investigating how terrestrial ectotherms respond to these new conditions provides an excellent opportunity to explore the broader impacts of a warming climate.

The challenge of maintaining optimal thermal conditions during embryonic development is central to reproductive success among many ectotherms, though overcoming this challenge is accomplished using different strategies. Oviparous (egg laying) ectotherms must ensure consistent thermal conditions for their eggs throughout the entire period of embryonic development by utilizing complex behaviors such as careful nest building, precise nest site selection, or by maintaining thermal conditions through monitored incubation of clutches (Du and Shine, 2014). Typically, these nest sites are abandoned by the parent shortly after egg deposition, allowing the nest itself to maintain environmental conditions conducive to proper embryonic development while the mother is free to forage and rebuild depleted energy reserves (Shine, 1988; Gans et al., 1996; Huang and Pike, 2013; Doody and Refsnider, 2022). In contrast, viviparous (live-bearing) ectotherms must alter their behavior while gravid to maintain optimal internal thermal conditions throughout gestation, even to the extent of risking their own survival to prioritize ideal thermal conditions for developing embryos (Seigel et al., 1987; Charland and Gregory, 1995; Gregory et al., 1999; Lorioux et al., 2013). This behavioral tradeoff serves to illustrate the importance of an organism’s thermal environment, and that maintaining proper temperature is vital for the survival and optimal development of offspring and therefore directly impacts the individual fitness of reproducing adults. Temperatures experienced during gestation are often observed to impact neonate size in viviparous squamates (Mathies and Andrews, 1997; Lourdais et al., 2004; Gao et al., 2010; King, 2022). Both greater neonate body length (Brown and Shine, 2004; Kissner and Weatherhead, 2005; Addis et al., 2017) and greater body mass (Jayne and Bennett, 1990; Bronikowski, 2000; Kissner and Weatherhead, 2005) are associated with significantly increased rates of survival to adulthood in viviparous snakes.

Changes to environmental thermal conditions during gestation have been observed to impact both physiology and behavior of viviparous ectotherms to alter characteristics such as maternal energy use and basking and foraging behavior (Lourdais et al., 2002; Ladyman et al., 2003; Wang et al., 2017), gestation length (Blanchard and Blanchard, 1940; Ji et al., 2006; Wang et al., 2017; King, 2022), litter size (Lourdais et al., 2004; Zhang et al., 2010; Tang et al., 2012), and body condition and viability of neonates (Lourdais et al., 2004; Wang et al., 2017; King, 2022). Although there is often agreement regarding the influence of warmer temperatures on gestation, neonate survival, and body condition, the specific effects observed differ between studies and depend on the biological system examined. Wang et al. (2017) reported that increased ambient temperature was associated with significantly shorter gestation length, increased neonate mass, and a decrease in general activity during gestation for Toad-headed lizards (*Phrynocephalus putajatia*). Beuchat (1988), found that increased environmental temperatures during gestation were associated with a decrease in gestation length in Yarrow’s spiny lizards (*Sceloporus jarrovi*), but this study also reported that sustained increased temperatures resulted in a detrimental effect on the body conditions of both mother and neonate. This latter general trend was further supported, as Tang et al. (2012) reported an increase in litter size and a decrease in gestation length associated with higher temperatures for Multi-ocellated racerunner lizards (*Eremias multiocellata*), but that these potentially beneficial effects were lost as temperatures continued to increase. Lourdais et al. (2004) demonstrated that increased ambient temperature resulted in shorter gestation length, increased embryo viability, and caused phenotypic changes, including changes to scale patterns, in neonate European asps (*Vipera aspis*) within a population living at the northern boundary of its range. Each of these studies demonstrate that, even among animals that are able to effectively behaviorally thermoregulate, increased ambient temperatures significantly impact both the behavior of gravid females and the resulting reproductive outcomes in viviparous species.

Red-sided garter snakes (*Thamnophis sirtalis parietalis)* are viviparous ectotherms and one of the most northerly living squamate species in North America (Aleksuik and Stewart, 1971). The focal population of this study is found in the Interlake region of Manitoba, Canada near the northern extreme of the species’ range which spans as far South as Northern Texas. The Interlake region is particularly well suited for studies examining the effects of temperature on ectothermic vertebrates, as it is characterized by major seasonal changes to the thermal landscape. Unsurprisingly, ectotherms living within this region are well adapted to this extreme environment and demonstrate highly coordinated annual behavioral cycles primarily driven by seasonal temperature changes (Aleksiuk and Stewart, 1971; Gregory, 1974; Lutterschmidt et al., 2006). Indeed, Red-sided garter snakes spend the majority of their life in a hibernation-like state of winter dormancy known as “brumation,” during which the entire population retreats into subterranean hibernacula for up to 8 months each year (Aleksiuk and Stewart, 1971; Costanzo, 1985). Brumation is a behavioral and physiological state characterized by extremely low levels of activity and greatly reduced metabolic rates in ectotherms. This state is driven by low temperatures and serves as an essential strategy for surviving extremely cold and resource-limited winter conditions (Gregory, 1982; Geiser, 2013). Red-sided garter snakes remain relatively inactive throughout brumation and do not feed for approximately 8-9 months each year (O’Donnell, 2004). Thus, they must rely solely upon stored energy to survive winter dormancy as well as the intense and energetically costly activities of mating and migration immediately following emergence from the hibernacula in early spring (Aleksiuk and Gregory, 1974; Aleksiuk, 1976). These energy stores must be obtained during an extremely brief summer feeding season lasting as little as 10 weeks, during which an individual must acquire all of the energy necessary for an entire year of growth, development, reproduction, and survival (Crews et al., 1987; O’Donnell et al., 2004). Moreover, the challenge of building energy reserves becomes even more difficult for reproducing females as this brief feeding period coincides with gestation – a period that is both behaviorally and energetically demanding. Unlike the southern populations, the environmental temperatures experienced by the northern populations are much lower relative to their optimal body temperatures, suggesting minimum ambient temperatures may play a large role in the behavior and reproductive outcomes for snakes undergoing gestation in this colder region (Gibson, 1977; Gibson and Falls, 1979; Arnold and Peterson, 2002). Thus, this limited window of time is critically important and has the potential to dramatically impact individual fitness, as a reproducing female must carefully balance activities and energy demands to simultaneously ensure survival while maximizing reproductive success.

Although climate change-induced warming has resulted in a global average temperature increase of approximately 0.6 °C over the last 60 years, the Interlake region has been disproportionately impacted, experiencing nearly triple the global rate with recorded temperatures showing an increase of 1.7 °C over the same period (Zhang et al., 2000; Houghton et al., 2001; Environment Canada, 2014; Bush and Lemmen, 2019). Global temperatures are predicted to continue to increase by as much as 5.3 °C by the end of the century, which is well beyond the findings of previous work where increases of as little as 2 °C significantly impacted gestation in viviparous ectotherms (Houghton et al., 2001; Kingsolver et al., 2013; Wang et al., 2017; Masson-Delmotte et al., 2021). With current regional warming levels so much higher than the global average, it is reasonably likely that future regional warming will be at the higher end of global climate change model projections. While increased temperature in this region will likely not result in thermal conditions that are significantly warmer than those currently experienced by this species at the southern end of their range, increases to minimum temperatures of this degree have a large potential to alter behavioral thermal regulation and reproductive outcomes for snakes at the colder northern end of their range. This is especially true for increases in nighttime temperatures, where even during summer they currently remain well below optimal gestation temperatures.

Currently, the daily mean environmental temperatures experienced by Red-sided garter snakes during the summer feeding and gestation period range from 13.8 °C in late September to 20.1 °C in July, with mean maximum daily temperatures 25 °C, and mean minimum temperatures 15.9 °C (https://en.climate-data.org/north-america/canada/manitoba/inwood-108217/). Previous research focused on the Red-sided garter snakes and other members of the genus *Thamnophis* have shown that both free-ranging and captive gravid females behaviorally maintain daytime body temperatures between 26 °C and 31 °C – a range that typically exceeds ambient temperatures (Gibson and Falls, 1979; Arnold and Peterson, 2002; Hughes et al., 2019). Though often overlooked, nighttime temperatures are rising faster than daytime temperatures, resulting in a potentially larger impact to minimum temperatures than maximum temperatures, especially for organisms living at the coldest edges of their range (Alexander et al., 2006; Rutschmann et al., 2023)

In the present study, we manipulated the thermal environment of gestating female Red-sided garter snakes by increasing minimum diurnal and nocturnal ambient temperatures by an ecologically relevant 5 °C and observed the impact of this increase on gestation length and the neonate and litter-wide traits at birth. We predicted an increased minimum ambient temperature of 5 °C would significantly reduce gestation length, with potential increases to neonate mass, body condition, and parturition success. However, we also predicted that if this thermal increase exceeds the beneficial thresholds of this population, potential individual gains would be lost resulting in a reduction in neonate body condition, reduced offspring per litter, increases to stillbirths and severe deformities (Beuchat, 1988; Tang et al., 2012).

## 2. Materials and methods

All experimental protocols presented were approved by the Oregon State University Institutional Animal Care and Use Committee (Animal Care and Use Protocol 5058), in accordance with the Association for the Assessment and Accreditation of Laboratory Animal Care. This research was conducted under the authority of Manitoba Wildlife Scientific Permit No. WB20333.

### Animal Collection

We collected free-ranging male and female Red-sided garter snakes (approximately 100 of each) from the natural population at over-wintering hibernacula near the town of Inwood, Manitoba, Canada during the spring mating season between 1 May and 4 May 2018. This site contains several communal hibernacula that collectively shelter a population estimated at approximately 35,000 snakes (Shine et al., 2006). Male snakes were selected and captured after being observed actively exhibiting courtship behavior. Non-mated females were selected and captured only if their initial emergence from the hibernaculum was observed, and they were being actively courted by males. To minimize the effects of body size variation on litter size, only females of the same approximate size were selected, with snout-vent length (SVL) and body masses of approx. 60 cm and 100 g, respectively. Due to the extremely large population and ease of collection, we were able to limit collection to this target size while easily maintaining a sufficient sample size. To confirm that the females had not yet mated that season, we visually inspected their cloaca for copulatory plugs upon collection. Copulatory plugs are large, gelatinous spermatophores deposited by a male inside of a female’s cloaca during copulation. Copulatory plugs are substantial in size, persist for several days, and represent an easily identifiable and reliable indication that a female has recently mated (Whittier et al., 1985; Friesen et al., 2013). Any female presenting a copulatory plug, or any indication of potentially having mated, was immediately released. All animals were collected by hand and placed in cloth bags for transportation to the Chatfield Research Station, Chatfield, Manitoba. Snakes were separated by sex and housed outdoors in specially designed 1 m^3^ nylon enclosures containing natural stones, hide boxes, and water dishes. Water was provided *ad libitum* for the duration of captivity, but food was not provided in the field or at the research station, as this species is aphageous during the spring mating season (Crews et al., 1987, O’Donnell et al., 2004). Capture, handling, and transportation are not likely to have impacted results, as previous studies have consistently shown that these activities do not adversely affect survival, reproduction, or courtship behavior (Friesen et al., 2013; Friesen et al., 2014).

### Mating Trials

To control mating conditions and timing, we used courtship arenas (opaque, cylindrical, nylon enclosures 0.5 m in diameter and 1 m tall) with heat lamps (above) and electric heating pads (below) to maintain homogenous thermal environments at ≥ 13.0 °C – optimal temperatures to stimulate vigorous courtship behavior. Female snakes were placed individually into arenas and paired with 5 randomly selected males. All snakes were allowed to freely behave under constant observation until successful copulation occurred. Successful copulation was confirmed by visual inspection and identification of a copulatory plug. Females with copulatory plugs were separated from the males and transferred to a 1 m^3^ outdoor nylon enclosure containing only mated females. Each male was allowed to mate only once. Males that successfully mated were removed from the remainder of the study to minimize any potential bias that may arise from allowing a male to sire more than a single litter of offspring. All males were marked dorsally with indelible ink to prevent recapture and released at the site of capture within 5 days of collection. All controlled matings took place within the same 48-hour period, concluding when 80 successful matings occurred. Similar experiments have reported that 23-64% of mated female Red-sided garter snakes give birth in a laboratory setting (Friesen et al., 2014; Blakemore 2017; Dayger et al., 2018), as such our sample size should result in 10-20 successful litters per treatment.

### Experimental Conditions and Data Collection

We transported the mated females (*n* = 80) by car to the laboratory facilities at Oregon State University. Animals were transported in cloth bags, housed in a thermally controlled environment (air temperatures between 20 °C and 22 °C), and given water via a damp sponge to prevent dehydration for the duration of the trip (< 30 h). Though females were all approximately the same size as a result of carefully targeted collection, further efforts were made to ensure any remaining size variation was accounted for between treatment and control groups. Female snakes were visually sorted by SVL, then matched into pairs based on both SVL and approximate body condition to minimize the impact of differences in energy reserves and maternal body size between treatments. These matched pairs were then randomly split between treatment and control conditions, from largest to smallest to ensure an even distribution of size ranges in each group of 40 snakes. Snakes were marked ventrally via scale clipping to indicate both the experimental condition and the relative size ranking within each treatment. Once assigned to a group, and for the remainder of the study, snakes were housed within microprocessor-controlled environmental chambers (Environmental Growth Chambers, Chagrin Falls, OH). The control group was housed within a chamber emulating the thermal environment found in this species’ natural range during the summer feeding period, which consists of a daytime temperature of 22 °C and a nighttime temperature of 15 °C. The treatment group was housed in an identical chamber set to maintain ambient temperatures increased by 5 °C, which consists of a daytime temperature of 27 °C and a nighttime temperature of 20 °C. Females were housed individually in 45 liter glass aquaria containing approximately 2.5cm of CareFresh™ paper fiber substrate, a water bowl, a hide box, a rock to facilitate shedding of skin, and a heat lamp situated at one end to produce a thermal gradient allowing behavioral thermoregulation within the constraints of the experiment. Maximum temperatures under the basking lamps were approximately 31 °C for control group and 40 °C for the treatment group. Minimum temperatures never dropped below the ambient temperature for each treatment (22 °C:15 °C or 27 °C:20 °C day:night cycle). Thus, this study tightly controlled the minimum ambient temperatures snakes were able to access during gestation while allowing snakes to freely thermoregulate and access higher temperatures if preferred, mimicking thermal opportunities experienced in the wild where minimum temperatures are reached at night but behavioral thermal regulation during the day allows for higher diurnal body temperatures. Photoperiods were identical for both treatments and emulated that of the summer feeding period in Manitoba (16 h:8 h light:dark cycle). Humidity was maintained at 65 % for both treatments. Water was provided *ad libitum*, and food (a mixture of earthworms and salmon fry supplemented with calcium) was provided once per week, during which time all snakes were allowed to eat to satiation. Snakes were maintained in these conditions for the duration of the experiment.

Each female was inspected daily to identify births, and offspring were collected within no more than 24 hours of birth. Dates of birth and number of offspring were recorded for each litter. Neonates were weighed and measured immediately after collection. Masses were recorded to the nearest 0.01 g and SVLs were recorded to the nearest 1 mm. Measurements were collected for every litter and for every neonate until the last female gave birth in late September (see Supplemental Table 1 for full data). All adult snakes were maintained in captivity until the following spring then returned to the location of capture. Offspring were maintained in the lab for further data collection facilitating additional studies aiming to characterize growth and development.

### Statistical methods

To assess the overall health of neonates, Scaled-mass index (SMI), a non-destructive method for estimating body condition in animals using mass and length data was calculated for each neonate (Peig and Green, 2009). Briefly, SMI was calculated from the following equation:

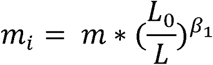

Where *m_i_*is the scaled-mass index, *m* is the individual mass, L_0_ is the mean SVL, L is the individual SVL, and β*_1_* is the β*_1_*coefficient derived from the following linear mixed-effects model (LMM):

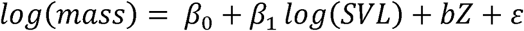

Where β*_0_* is the intercept, β*_1_* is the coefficient for the fixed effect of log(SVL), *b* is the coefficient for the random effect of Z (mother), and ε is the error term. LMM and SMI were calculated for each treatment condition. Using an LMM allows us to account for interlitter variation, such as minor differences in maternal physiology, behavior, and environmental conditions associated with each litter by treating the mother as a random effect. Additionally, LMMs are robust to violations of normality and independence, allowing us to make full use of the dataset without limiting analyses to mean litter values for each trait, as well as enabling the use of non-transformed data to generate results that can be directly interpreted (Bates et al., 2015; Schielzeth et al., 2020).

To evaluate the effect of treatment on reproductive outcomes, a linear mixed effects model (LMM) was used to evaluate the effect of treatment (treatment or control) on traits including neonate mass (g), SVL (cm), SMI, as well as gestation length (days) using the following:

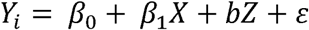

Where *Y_i_* is trait of interest, β*_0_* is the intercept, β*_1_* is the coefficient for the fixed effect of X (treatment), *b* is the coefficient for the random effect of Z (mother), and ε is the error term. A full model that also included relative maternal size as fixed effect, as well as a reduced model that accounted for the random effects of mother but did not include the effect of treatment were compared using the Akaike information criterion (AIC), however the intermediate model above resulted in the best fit and is used throughout (Supplemental Table 2). All LMM analyses were conducted using the lme4 package (Bates et al., 2015) in R (R Core Team, 2023). All figures created using the ggplot2 package in R (Wickham, 2016).

To explore the potential treatment-based differences in the relationships between traits measured, an LMM that is similar to the one above was employed, however the X variable instead represented one of the traits. Maternal identity was still included as a random effect to account for interlitter variation. For each model, the marginal R^2^ was calculated using the ‘r.squaredGLMM’ function from the MuMIn package in R (Barton and Barton, 2015). Traits compared this way include neonate mass, SVL, SMI, gestation length, and litter size.

Additionally, interactions between traits and treatment conditions were evaluated for the traits neonate mass, neonate SVL, and neonate SMI using an expanded LMM that included the interactions between neonate SVL, neonate mass, gestation length, litter size, and treatment condition, with maternal identity held as a random effect. Relationships were then visualized by plotting two traits against each other and fitting an LMM to each treatment using the ggplot2 package in R.

To further evaluate the effect of treatment on reproductive outcomes we used a linear model (LM) to compare litter outcomes between treatments (See Supplemental Table 3 for litter wide traits). Treatment (treatment or control) was treated as a fixed effect on traits including total litter mass (g), average neonate mass (g), average neonate SVL (cm), average neonate SMI, litter size, number of stillborn, and number of deformed neonates per litter, using the following equation:

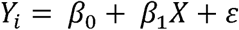

Where *Y_i_* is the trait of interest for each litter, β*_0_* is the intercept, β*_1_* is the coefficient for the predictor X (treatment), and ε is the error term. All LM analyses were conducted using the stats package (version 4.3.1) in R. All figures created using the ggplot2 package in R.

Additional analyses were conducted to explore the relationship between relative maternal size and those same traits, however, due to the ranked nature of the relative maternal size, Kendall’s Tau correlation was calculated using the ggpubr (version 0.6.0) package in R. All correlation figures were visualized using ggplot2.

A two proportions z test with continuity correction was used to evaluate the proportion of females that gave birth out of the total number of mated females for each treatment using the prop.test function of the stats package in R.

## 3. Results

A total of 26 mated females reached parturition, with 12 females from the treatment and 14 from the control group successfully giving birth. The proportion of mated females that reached parturition was well within the reported range for a laboratory setting, with 35% of control and 30% of treatment snakes successfully giving birth. When compared, there was no significant difference in birthrate between treatments (z^2^ = 0.06, p > 0.8, n_treatment_ = 12, n_control_ = 14) (Figure 1L).

**Figure 1.**
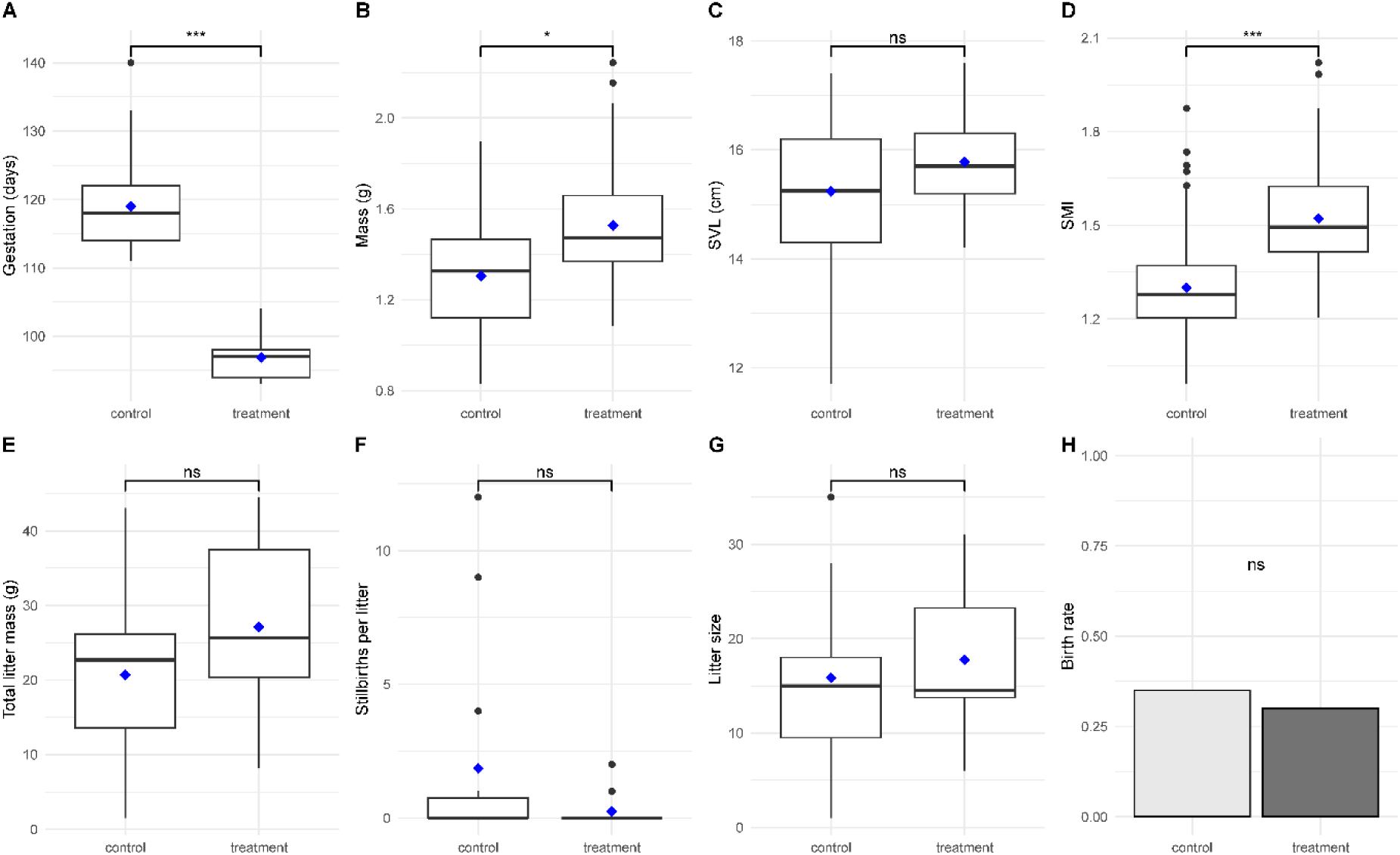
Differences in reproductive outcomes for snakes held at different minimum temperatures during gestation. Treatment condition represents snakes housed at a constant 5 °C higher minimum temperature relative to control for the duration of gestation. Boxes indicate the median (horizontal line in box), the first (Q1) and third (Q3) quartiles represent the values below the 25^th^ percentile or above the 75^th^ percentile respectively (bottom and top of box) and mean (blue diamond), whiskers represent the largest or smallest value within 1.5 x interquartile range (IQR). Barplot indicates the proportion of mated females that reached parturition for each condition. Statistical results from linear mixed-effects models on individual neonate outcomes (A-D), linear models on litter-wide outcomes (E-G), and a two proportions z-test (L). Gestation length was significantly reduced (A) and neonate mass and body condition (SMI) were significantly higher in the treatment condition (B and D respectively). There was no difference in neonate snout-vent length (SVL) between conditions (C). There was no significant effect caused by treatment at the litter level for total litter mass (E), number of stillbirths (F), or litter size (G). There was no significant difference in the birthrates (L) between treatments. Statistical significance is indicated as: ***p < 0.001, **p < 0.01, *p < 0.05, and ns (p > 0.05).

Neonate mass, SMI, and gestation length were all found to be significantly affected by the treatment condition (p < 0.05), with no difference present in SVL between treatments (p > 0.5) (Figure 1A-D and Table 1). The effect of treatment increased both neonate mass and SMI, with an estimated increase of 0.19 g and 0.23 respectively, and decreased gestation length by an estimated 24 days. This shows that not only did the treatment condition reduce gestation length by approximately 20%, but also increased neonate mass by approximately 14% while maintaining neonate SVL.

**Table 1.**
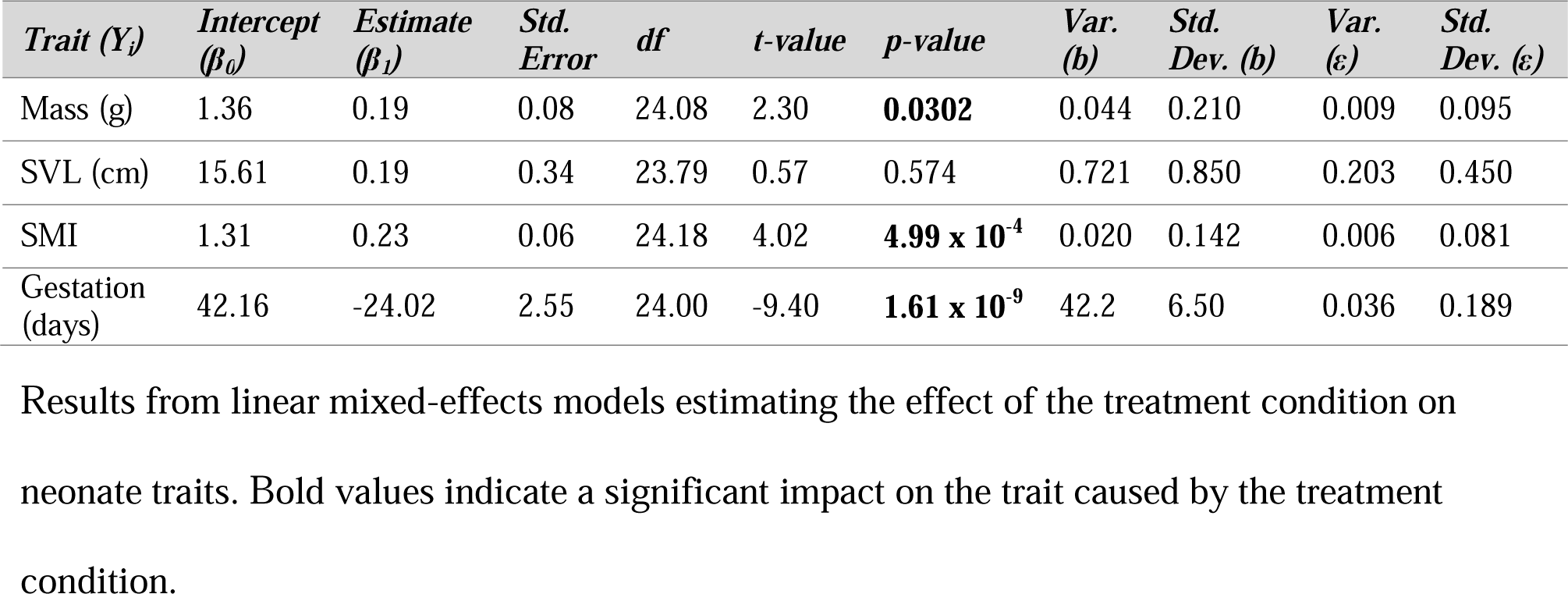
The effect of treatment on neonate traits at birth.

| *Trait (Y<sub>i</sub>)* | *Intercept (β<sub>0</sub>)* | *Estimate (β<sub>1</sub>)* | *Std. Error* | *df* | *t-value* | *p-value* | *Var. (b)* | *Std. Dev. (b)* | *Var. (ε)* | *Std. Dev. (ε)* |
| --- | --- | --- | --- | --- | --- | --- | --- | --- | --- | --- |
| Mass (g) | 1.36 | 0.19 | 0.08 | 24.08 | 2.30 | **0.0302** | 0.044 | 0.210 | 0.009 | 0.095 |
| SVL (cm) | 15.61 | 0.19 | 0.34 | 23.79 | 0.57 | 0.574 | 0.721 | 0.850 | 0.203 | 0.450 |
| SMI | 1.31 | 0.23 | 0.06 | 24.18 | 4.02 | **4.99 x 10<sup>-4</sup>** | 0.020 | 0.142 | 0.006 | 0.081 |
| Gestation (days) | 42.16 | -24.02 | 2.55 | 24.00 | -9.40 | **1.61 x 10<sup>-9</sup>** | 42.2 | 6.50 | 0.036 | 0.189 |

Results from linear mixed-effects models estimating the effect of the treatment condition on neonate traits. Bold values indicate a significant impact on the trait caused by the treatment condition.

Of the litter-wide outcomes including total litter mass (g), mean neonate mass per litter (g), mean neonate SVL per litter (cm), mean neonate SMI per litter, litter size, number of stillbirths per litter, and number of significant developmental deformities per litter, only mean neonate mass per litter and mean neonate SMI per litter were significantly different between treatments (p < 0.05), with the estimated effect of treatment consistent with those found in the LMM results (0.19 g and 0.23 respectively) (Figure 1E-G, Supplemental Figure 1, and Table 2).

**Table 2.**
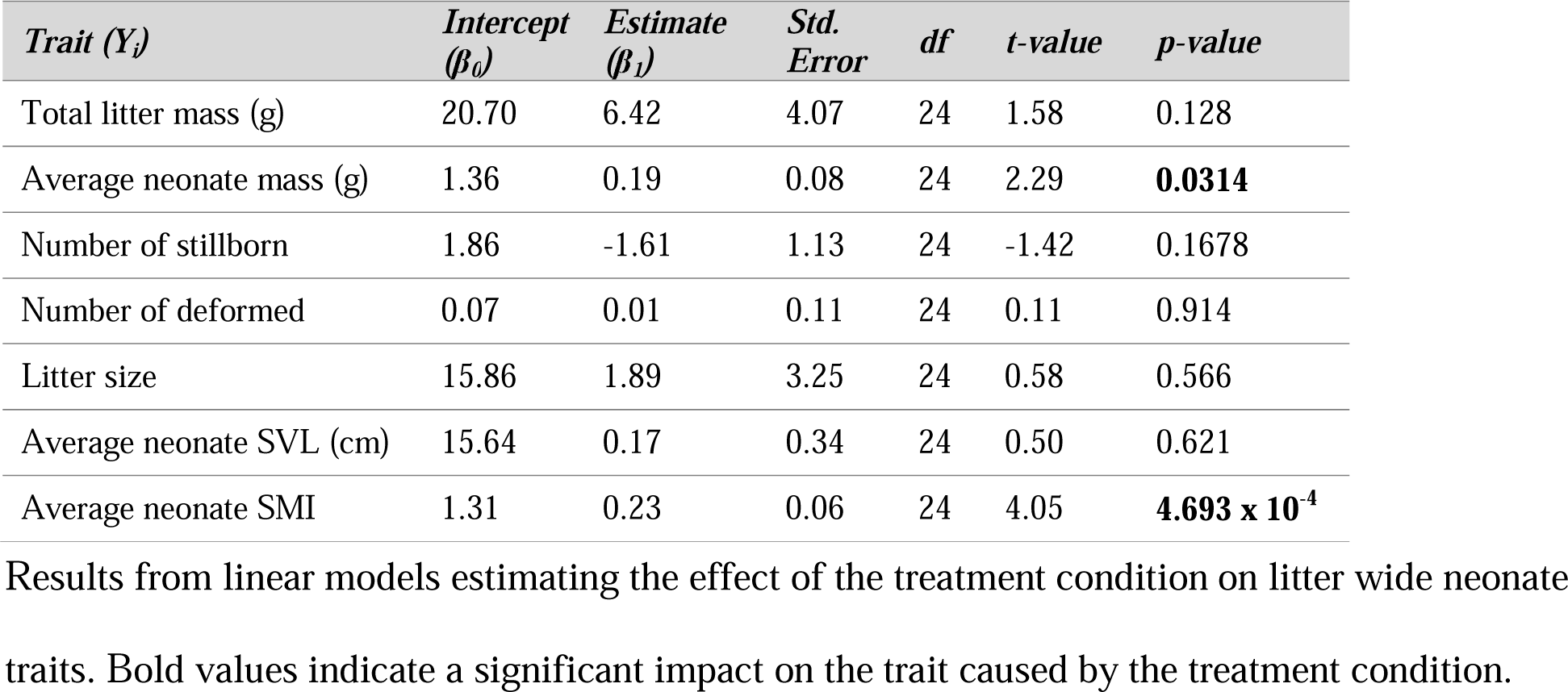
The effect of treatment on traits of neonates at birth by litter.

A weak positive relationship was found between neonate SVL and neonate mass for treatment (F_[1, 209]_ = 8.86, p < 0.001, R^2^ = 0.18, n = 213) and a positive relationship for control (F_[1, 220]_ = 214, p < 0.001, R^2^ = 0.41, n = 222) (Figure 2B). When plotted against SMI, neonate SVL and neonate mass illustrate detail in how SMI differed between treatment groups. Neonate mass had a strong positive relationship with neonate SMI for treatment (F_[1, 95]_ = 25, p < 0.001, R^2^ = 0.89, n =213), and a similar relationship for control (F_[1, 218]_ = 21.2, p < 0.001, R^2^ = 0.73, n = 222) (Figure 2C). Neonate SVL showed no relationship with neonate SMI for control (F_[1, 214]_ = −0.135, p = 0.893, R^2^ < 0.001, n = 222), or for treatment (F_[1, 210]_ = −0.162, p = 0.871, R^2^ < 0.001, n = 213), showing both that SMI is behaving as expected, and that there is a consistent shift to a higher SMI in the treatment condition regardless of SVL (Figure 2A). The treatment group showed mass that skewed higher at larger SVL, suggesting that while there was no difference in SVL between treatments (Figure 1C), neonates with a higher SVL were able to build more mass in the treatment group compared to the control group, where neonates had appear to have had difficulty building mass for longer individuals.

**Figure 2.**
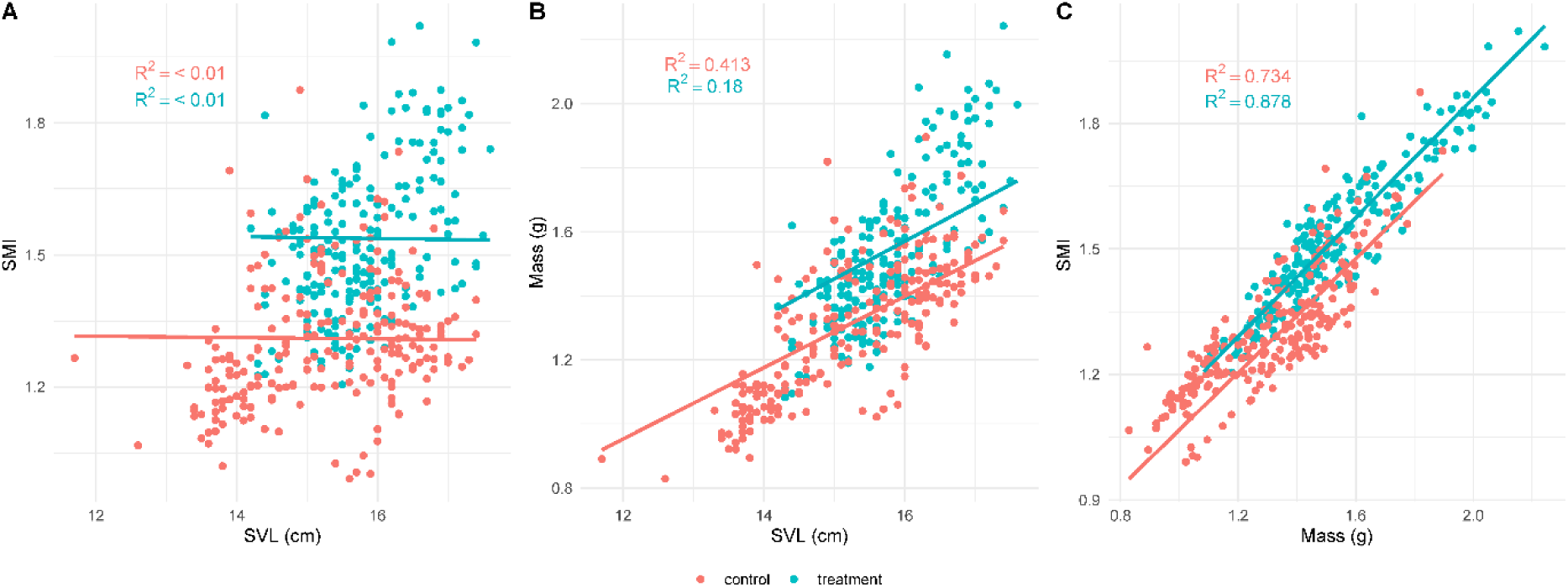
Relationship between neonate traits. Lines indicate liner mixed-effects regression model (LMM) fit, color coded by. Marginal R^2^ is displayed for each LMM, by treatment. Neonate snout-vent length (SVL) plotted against scaled-mass index (SMI) illustrates the difference in SMI between treatment and control groups, with the treatment group showing higher SMI (A). A positive relationship between neonate SVL and neonate mass for both treatment conditions (B). Neonate mass plotted against SMI shows that the relationship between the two is similar for both treatment conditions, but shifted higher for the treatment conditon (C). See Supplemental Table 4 for full linear mixed-effects model results.

Litter size had little to no relationship with, neonate SVL, neonate mass or neonate SMI for the treatment group, but a strong slightly negative relationship was found between litter size and gestation length (F_[1, 0.006]_ = −10.3, p = 0.969, R^2^ = 0.901, n = 213). A negative relationship was found between litter size and neonate mass (F_[1, 12.9]_ = −2.27, p = 0.47, R^2^ = 0.23, n = 222), neonate SVL (F_[1, 13.1]_ = −3.89, p < 0.01, R^2^ = 0.45, n = 222), and gestation length (F_[1, 12]_ = −2.75, p <0.05, R^2^ = 0.36, n = 222) for the control group (Supplemental Figure 2 and Supplemental Table 4). This suggests that cost associated with larger litters for the treatment group was generally very small, but controls with larger litters had moderate reductions to neonate mass and SVL as well as shorter gestation length. No relationship between gestation length and neonate traits were found for either treatment or control groups (all p > 0.1 and R^2^ < 0.1) (Supplemental Figure 3 and Supplemental Table 4). When looking at effect of the interaction between traits and treatment conditions on neonate mass, neonate SVL, and neonate SMI, a number of interactions were significant, with 17/21 significant interactions including the term for treatment condition (p < 0.05), further showing the strong impact of the treatment condition (Supplemental Table 5).

When comparing relative maternal size to neonate traits, we see several relationships that are significant, however all of these relationships show a very weak effect (Supplemental figure 4 and Supplemental Table 6). We believe that this shows that while maternal physiology plays a large role in neonate physiology, the weakness of the relationships in our data reflects our efforts to control for differences in maternal size during our initial sample selection.

All code used for the statistical analysis above and for all figures presented can be found at https://github.com/hubertdl/Tsp_Gestation_and_ClimateChange.

## 4. Discussion

Our observations of gestation times accurately agreed with our initial prediction that a 5 °C increase in minimum temperature during gestation would result in a reduction in gestation length. Indeed, we found that gestation lengths were significantly reduced in the treatment condition, such that the extent of the resulting ranges of birth dates did not overlap between the treatment and control groups. The average gestation length for the treatment was 24 days shorter than that of the control group representing a 20 % decrease in average gestation length. Shorter gestation lengths may increase the female’s survival as gravidity imposes energetic costs as well as reduced locomotory and foraging capabilities and requires increased amounts of time engaged in active thermoregulation – all of which may ultimately result in an increased exposure and risk of predation for the gravid female (Seigel et al., 1987; Charland and Gregory, 1995; Gregory et al., 1999). As these costs in Red-sided garter snakes are imposed simultaneously with the already extremely limited opportunities for summer foraging, a reduction in gestation length would allow for both increased amount of time to forage each year and more frequent prey capture during this period ultimately resulting in increased opportunity to replenish depleted energy stores after giving birth. Additionally, it has been shown that higher temperatures increase the ability to build mass during the summer season in adult females, allowing them to further make use of potentially expanded foraging in these hotter conditions to accelerate postpartum recovery (Halliday and Blouin-Demers, 2018).

However, it must be considered that any increase in foraging opportunity may be counterbalanced by potential decreases in prey availability, as increased temperatures will undoubtedly affect the whole ecosystem in ways that are difficult to predict. Additionally, climate change results in more frequent and extreme temperature swings, which will likely result in an increased occurrence of conditions that are outside the thermal range needed for normal activity, potentially reducing the time available for foraging per day (Masson-Delmotte et al., 2021). Despite this uncertainty, any increase in the time available for foraging will potentially increase the ability to overcome the challenges that result from reductions to suitable foraging conditions and prey availability. While warming conditions will likely alter seasonal timing, resulting in some extension to the spring and fall that are available for activity, winter brumation conditions are unlikely to change in a meaningful way under current climate change predictions, as current mean temperatures are well below freezing with sustained snow coverage from late October to mid-April each year (https://en.climate-data.org/north-america/canada/manitoba/inwood-108217/). While the longest period of brumation is unlikely to change, even a marginal shift that reduces brumation duration at the margins will add an additional variable that can alter the impact of a reduced gestation length.

We found that, along with the reduced gestation length, the treatment condition also resulted in both increased neonate mass and SMI at birth but did not affect SVL. This was most pronounced when comparing neonates with larger SVL measurements, suggesting that control neonates had a harder time building mass for longer neonates. Generally, larger neonate mass and body condition at birth are associated with both improved short term and lifetime survival in snakes (Bronikowski, 2000; Brown and Shine, 2004; Kissner and Weatherhead, 2005; Addis et al., 2017). Based on the mass and SMI results alone, the offspring gestated in warmer conditions may have increased overall fitness resulting in better parental reproductive outcomes. However, there are potential trade-offs associated with accelerated development that can have long-term consequences, such as reductions in longevity, stress resistance, and future fecundity, any of which might overshadow the benefits gained by their larger size at birth (Chippindale et al., 2004; Shalev and Belsky, 2016). Long term studies tracking lifetime outcomes would be necessary to fully determine if the potential short-term survival benefits are outweighed by long-term costs of accelerated development, however it takes up to 3 years to reach sexual maturity, and a lifespan in the wild that can exceed 10 years, making this extremely logistically difficult to conduct, and far outside the scope of this study (Gregory, 1977; Bronikowski, 2008; Rollings et al., 2017).

Interestingly we observed no differences in birth rates, average litter sizes, total litter mass, the number of stillborn or severely deformed neonates between treatments. It has been shown that increased temperatures during gestation can reach a threshold, after which improved outcomes observed under moderate temperature increases begin to be overtaken by detrimental outcomes for developing offspring or gestating mothers, such as reductions to body conditions of the neonates, reduced litter sizes, and increased stillbirths (Beuchat, 1988; Yan et al., 2011; Tang et al., 2012). During this study, by artificially increasing the minimum ambient temperatures available to mothers with developing offspring, we prevented them from thermoregulating at a temperature below such a threshold. This serves to simulate and evaluate the effect of general warming in an environment with thermal conditions that are lower than preferred body temperatures. Under these conditions, our observations suggest improved reproductive outcomes at the time of birth, with no directly observed negative results associated with the treatment condition. Thus, we conclude that an increase of 5 °C to the minimum ambient temperatures during gestation likely did not exceed the threshold at which detrimental outcomes overcome the benefits gained. Generally, this suggests that there is a potential for additional beneficial effects from an increased minimum ambient temperature beyond 5 °C during gestation, as the benefits presented here suggest that they have not yet reached their detrimental thermal threshold. Though it is possible that some benefits may be present with a more moderate increase in temperature, such as increased litter size or birthrates, and may have already been lost to some degree at this higher temperature. Further research exploring a wider range of thermal conditions during gestation would be interesting, as this would expand our perspective allowing for increased ability to predict the effects of future increases in ambient temperatures and would potentially pinpoint the maximum ambient temperature threshold at which beneficial results began to reverse and detrimental impacts begin to arise.

While our experimental design increased both minimum diurnal and minimum nocturnal temperatures by 5 °C, we believe that the treatment effect was driven primarily by the increased nocturnal minimum temperature. In fact, the large thermal differences experienced by populations at different geographic regions are likely overcome using behavioral plasticity, as they exhibit the same diurnal thermal preference regardless of large differences in latitude and climate (Rosen, 1991). During the day, gravid females were allowed to use basking to behaviorally thermoregulate, mimicking behavior exhibited in natural settings, however the minimum temperature experienced during the night cannot be overcome using this strategy. This should result in relatively little difference in body temperature during the day but would allow for body temperatures to remain closer to optimal gestation temperatures for the treatment group at night. Because higher maternal body temperatures accelerate embryonic development, a higher minimum temperature experienced during the night has the potential to increase the rate of development at the time it is the slowest (Beuchat, 1988; Lorioux et al., 2013). Further studies looking specifically at increased nocturnal temperatures in this system would be valuable in deciphering the impact of this increasingly relevant change in thermal conditions (Rutschmann et al., 2023)

This study did not identify that critical threshold, but these results suggest interesting implications for future warming associated with continued climate change. The response variables examined here were limited and did not directly address or account for other phenotypic affects that may have arisen from increased temperatures during gestation. However, evidence suggests that many other developmental processes may be influenced by altered thermal conditions, and increased temperatures may result in changes to offspring phenotype such as differences in scale and bone development (Arnold and Peterson, 2002; Lourdais et al., 2004). Additionally, the impacts of temperature during gestation on offspring traits such as the aforementioned changes to morphology, body size, and body condition are frequently associated with both beneficial and detrimental long-lasting impacts on survival after birth (Bronikowski, 2000; Brown and Shine, 2004; Kissner and Weatherhead, 2005; Noble et al., 2018).

Current climate change models predict future global average temperature increases of up to 5.3 °C by the end of the century (Houghton et al., 2001; Masson-Delmotte et al., 2021). Thus, the results of this experiment show that an increase in average temperature of this degree, specifically during the brief yet crucial opportunity for feeding and gestation, is not likely to cause direct detrimental effects on the reproductive outcomes measured in this study for northern populations of Red-sided garter snakes and have the potential to improve neonate fitness. In fact, because this species spans much of North America and different populations experience a wide range of thermal conditions, the populations at the colder high latitudes are likely able to survive much warmer conditions similar to those at the southern edge of their range (Fitch, 1965; Aleksiuk and Stewart 1971).

However, the extended impacts of increasing temperature are likely to result in other adverse effects on this population. Thermal stress caused by exposure to increased temperature extremes (acute high or low temperatures) is known to result in a number of detrimental effects including misaligned behavioral timing, altered development, reduced or altered activity, and higher mortality rates (Houghton et al., 2001; Jentsch et al., 2009; Paaijmans et al., 2013; Stott, 2016). The greater thermal instability associated with climate change conditions may also lead to thermally induced alterations to energy use and metabolic rate (Friesen et al., 2015; Wilson, 2020; Holden et al., 2021). The combination of physiological and environmental changes due to increased environmental temperatures are also known to increase pathogen exposure and increase individual susceptibility to infection (Walther et al., 2001; Harvell et al., 2009; Gallana et al., 2013; Nnadi and Carter, 2021). Additionally, many species have been observed to alter seasonal migration patterns and shift territorial ranges to mitigate changes to their traditional ecosystems, ultimately resulting in disruption of ecosystems arising from novel interspecific competition and alterations to the availability of prey species or trophic effects (Parmesan, 2006; Gilman et al., 2010; Urban et al., 2012; Salinas Ramos et al., 2020). Although we observed effects of increased temperatures that appeared likely beneficial to offspring, it is possible that that the lifetime reproductive potential of mothers would in fact be reduced as a result of long-term exposure to increased temperatures or repeated bouts of rapid gestation. Thus, future research regarding changes to ambient temperatures during gestation may benefit from examining both the immediate and long-term health impacts of mothers, such as impacts on body condition, future fecundity, parasite load, and subsequent winter survival, in addition to the long-term survival and impacts on the physiology and long-term reproductive potential of offspring.

Higher temperatures caused by future warming have the potential to change the shape of this population, resulting in offspring that not only develop faster, but are born larger. In the present study, we observed no apparent tradeoff or cost to the overall number of offspring in each litter. This has the potential to dramatically increase the survival rates of future generations, while increasing the time each year spent foraging for both the mothers and the resulting offspring, which may serve to ameliorate potential decreased foraging success caused by reduced prey availability or suitable foraging temperatures (Seigel et al., 1987; Charland and Gregory, 1995; Brown and Shine, 2004; Kissner and Weatherhead, 2005).

The myriad impacts of climate change on an ecosystem are overwhelmingly complicated and interconnected. Thus, it is exceedingly difficult to develop models that accurately predict comprehensive, species-specific outcomes in response to altered thermal landscapes. The present study examined the effect of altered minimum temperatures during gestation within a single large population and thus represents only a single piece of the complex overall puzzle. As global temperatures continue to rise, additional research examining the effects of increasing temperature on the behavior and physiology of individual species, as well as studies encompassing entire ecosystems, will provide the invaluable insight necessary to further our understanding of how temperature affects ecosystems and, more importantly, will enhance our ability to predict the impacts of future climate change on the global ecosystem.

## 5. Conclusions

Our research demonstrated that, even under a worst-case scenario climate change model prediction, increased minimum temperatures may lead to improved reproductive outcomes for terrestrial ectotherms living at the northern edge of their range. Reduced gestation length has the potential to facilitate increased fitness for both mother and offspring, and while further work is needed to determine the long-term consequences of this altered gestation, this study found no observed cost to reproductive outcomes.

