## Supplemental Figures 1-4 for "Increased offspring size and reduced gestation length in an ectothermic vertebrate under a worst-case climate change scenario"

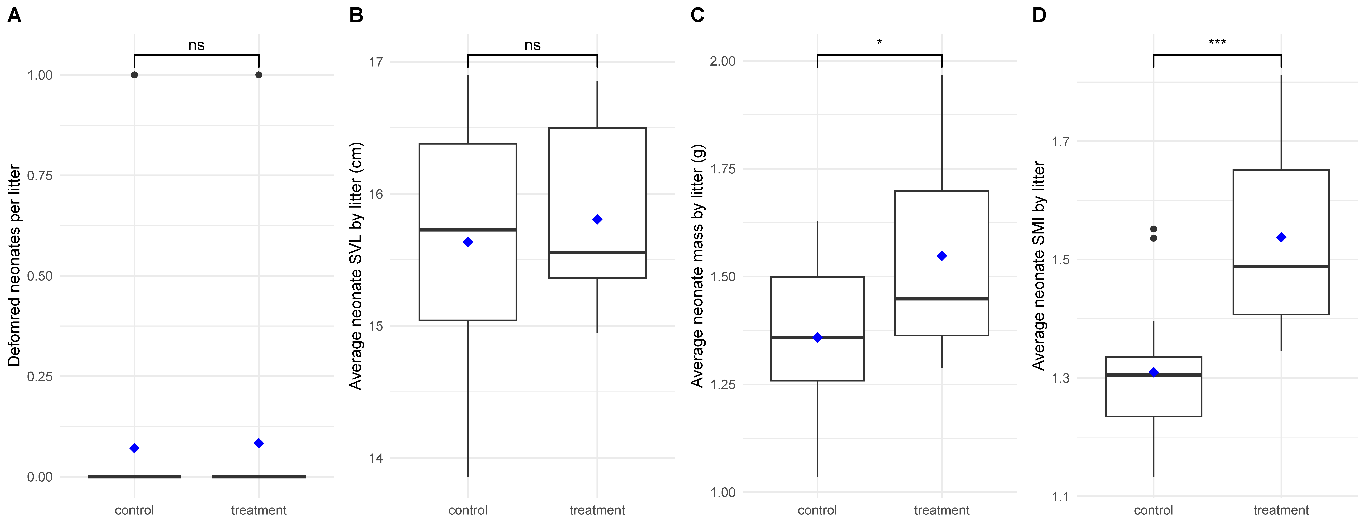

Supplimental figure 1. Differences in reproductive outcomes for snakes held at different minimum temperatures during gestation. Treatment condition represents snakes housed at a constant 5 ˚C higher minimum temperature relative to control for the duration of gestation. Boxes indicate the median (horizontal line in box), the first (Q1) and third (Q3) quartiles represent the values below the 25^th^ percentile or above the 75^th^ percentile respectively (bottom and top of box) and mean (blue diamond), whiskers represent the largest or smallest value within 1.5 x interquartile range (IQR). There was no difference in the number of deformed neonates per litter (A) or the average neonate SVL per litter (B). Both Average neonate mass per litter (C) and average neonate SMI per litter (D) were significantly higher in the treatment group when compared to the litter averages for the control group.

Statistical significance is indicated as: ***p < 0.001, **p < 0.01, *p < 0.05, and ns (p > 0.05).

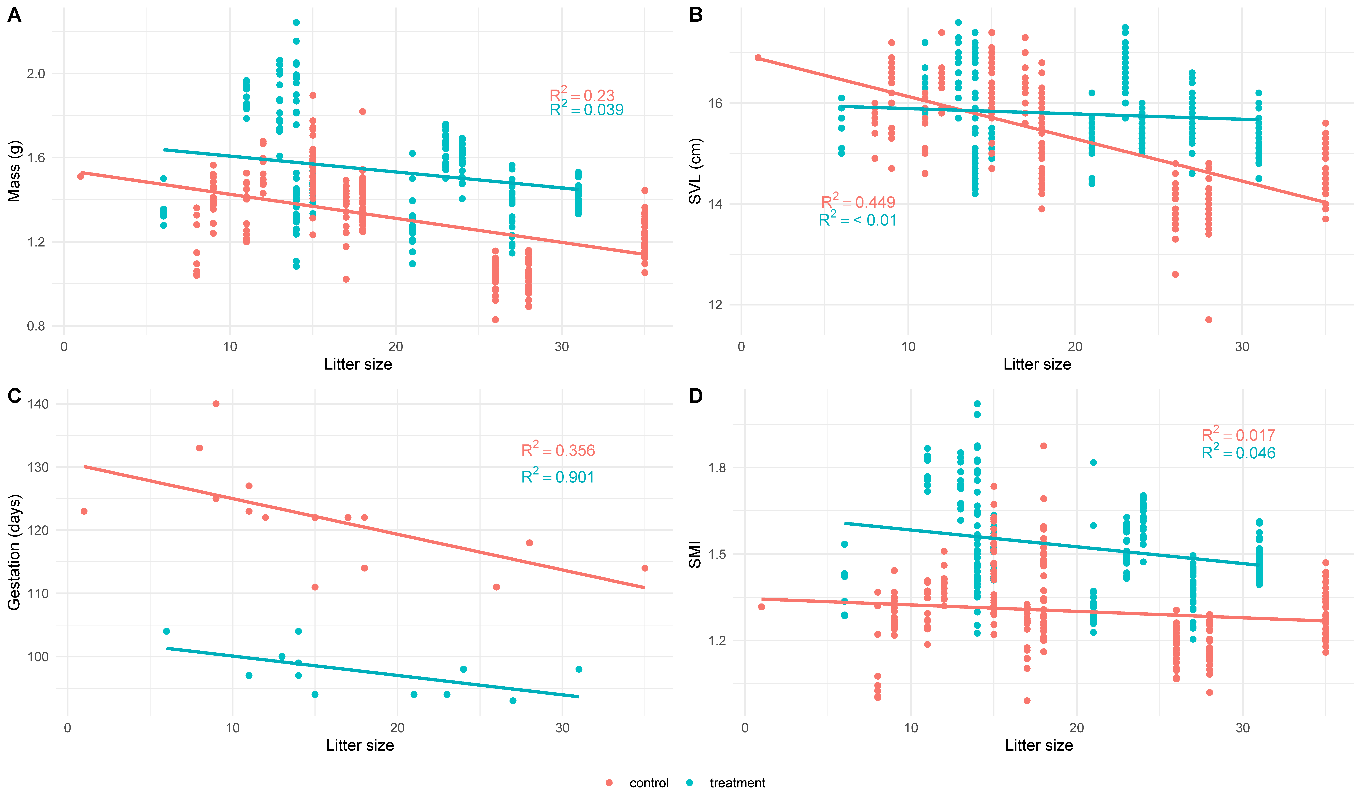

Supplimental figure 2. Relationship between neonate traits and litter size. Lines indicate liner mixed-effects model regression (LMM) fit, color coded by treatment. Marginal R^2^ is displayed for each model, indicating the strength of the explanitory power of litter size on the trait depicted. A negative relatioship between litter size and neonate mass (A), neonate SVL (B), and gestation length (C) is shown for the control group. There was a stron negative relationship between litter size and gestation length in the treatment group (C). Litter size had a very weak to no relationship with SMI in both treatments (D). See Supplemental Table 4 for full statistical results.

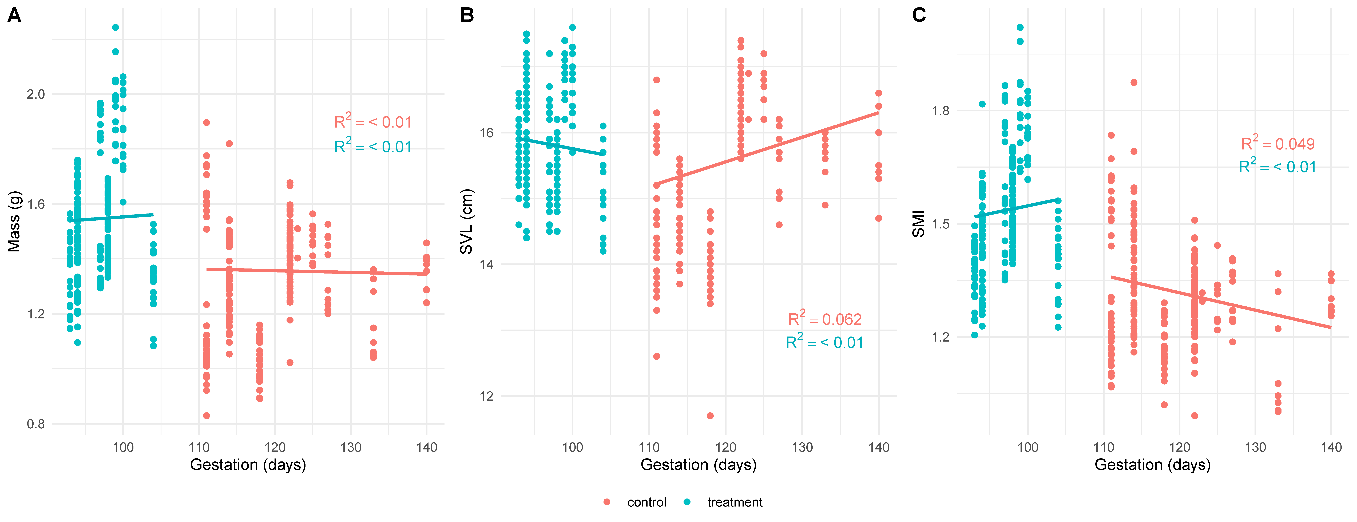

Supplemental figure 3. Relationship between neonate traits and gestation length in days. Lines indicate liner mixed-effects regression (LMM) fit, color coded by treatment. Marginal R^2^ is displayed for each model, indicating the strength of the explanitory power of gestation length on the trait depicted. Very weak to no relationship was found between gestation length and neonate mass (A), neonate SVL (B), or neonate SMI (C) for either treatment. (See Supplemental Table 4 for full LMM statistical results).

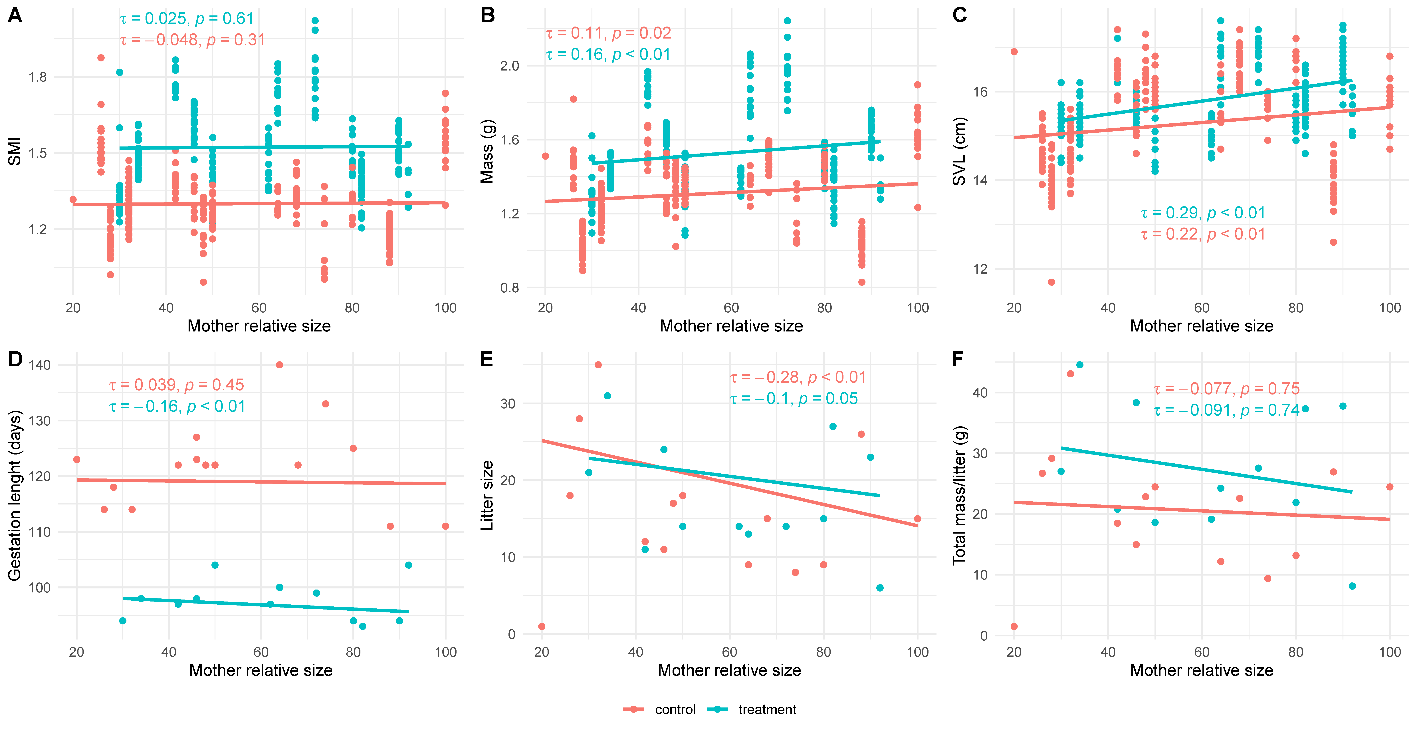

Supplemental figure 4. Relationship between neonate traits and relative maternal size. Lines indicate liner regression model fits, color coded by treatment. Tau and p-value are displayed for each model, indicating the results from a Kendell’s Tau correlation test. There was a very weak negative relationship between relative maternal size and SMI for both groups (A). There was a weakly positive relationship between relative maternal size and mass (B) as well as SVL (C) for both treatment conditons. There was a weakly negative relationship between relative maternal size and genstation length for the treatment conditon (D). Litter size had a weakly neagitve relationship with relative maternal size in both treatmetn conditions (E). No relationship was found between relative maternal size and total litter mass (F). See Supplemental Table 6 for full statistical results.
